# Mouse MRI shows brain areas larger in males emerge earlier than those larger in females

**DOI:** 10.1101/172841

**Authors:** Lily R. Qiu, Darren J. Fernandes, Kamila U. Szulc, Jun Dazai, Brian J. Nieman, Daniel H. Turnbull, Mark R. Palmert, Jason P. Lerch

**Author notes:** First authors who contributed equally (Lily R. Qiu, Darren J. Fernandes). Senior authors who contributed equally (Mark R. Palmert, Jason P. Lerch).

## Abstract

Sex differences exist in behaviours, disease and neuropsychiatric disorders. Sexual dimorphisms however, have yet to be studied across the whole brain and across a comprehensive time course of postnatal development. We used manganese-enhanced MRI (MEMRI) to longitudinally image male and female C57BL/6J mice across 9 time points, beginning at postnatal day 3. We recapitulated findings on canonically dimorphic areas, demonstrating the ability of MEMRI to study neuroanatomical sex differences. We discovered, upon whole-brain volume correction, that neuroanatomical regions larger in males develop early in life, while regions larger in females develop in peripubertal life. Furthermore, we found groups of areas with shared sexually dimorphic developmental trajectories that reflect behavioural and functional networks, and expression of genes involved with sex processes. Our results demonstrate the ability of MEMRI to reveal comprehensive developmental differences between male and female brains, which will improve our understanding of sex-specific predispositions to various neuropsychiatric disorders.

---

Sex differences in the brain are pervasive as demonstrated by differences in a wide range of processes including pain [1], learning and memory [2] and language [3]. Notably, there are robust sex differences in the prevalence, age of onset, and course of various psychiatric disorders. Males have a predisposition for disorders that have early onset during childhood, which include autism spectrum disorders, attention deficit disorders and Tourette syndrome. Females have a predisposition for disorders that have later onset, during adolescence and early adulthood, which include major depressive disorders, anxiety disorders and eating disorders [4]. Key to understanding these sex-specific vulnerabilities and predispositions is a better understanding of the normal development of sex differences in the brain.

Noninvasive image acquisition using MRI enables repeated scanning of the same individual to study longitudinal development. This is important because of the temporal nature of sex differences in the brain [5], behaviours, and risk factors for psychiatric disorders. Mesoscopic anatomy is also translatable between model organisms, such as mice, and humans due to the structural homology of their brains.

Anatomical sex differences have been studied in both rodents and humans. Histological studies in rodents have revealed some of the most well-known sex differences in the brain, such as the bed nucleus of the stria terminalis (BNST) [6], the medial nucleus of the amygdala (MeA) [6], and the medial preoptic nucleus of the hypothalamus (MPON) [7]. Due to the rodent’s accelerated lifespan, the sexually dimorphic development of these areas has been well-characterized beginning from neonatal life [8] [9] [10]. Using MRI, several anatomical sex differences have been found in the human brain as well. Whole brain size [11], the putamen, globus pallidus, basal ganglia, amygdala and hypothalamus are larger in males [11] [12] [13], while the caudate, thalamus and hippocampus are larger in females [12] [13] [14] Many of these sex differences arise during a critical period of sexual differentiation that occurs during neonatal life. The presence or absence of hormones in this developmental window organize the structure of the brain [15]. These sex differences are also subject to change throughout life as activational effects of hormones present during puberty and onwards can modulate neuroanatomy in males and females [16]. Additionally, sex chromosomes themselves influence sex differences in brain anatomy [17].

Due to the nature of sex differences in the brain and how they develop, there are shortcomings in the methods for investigating neuroanatomy in both rodents and humans. The accelerated life spans of rodents and their availability for histology renders them useful for investigating the development of specific nuclei; however, small nuclei are often examined in isolation, therefore neglecting others. In humans, MRI allows investigation of the whole brain, but lacks the ability to detect smaller sexually dimorphic nuclei. Furthermore, the long lifespan of humans renders studying development over a comprehensive period difficult. Most studies that investigate sex differences in the human brain are either cross-sectional, or examine sex differences on a relatively short developmental time scale. A recent meta-analysis examining sex differences in human brain structure across life determined that very few, if any, studies investigate sex differences during infancy and early childhood (0-6 years of age) [13], emphasizing the need to examine the neonatal window of development that encompasses a critical period of sexual differentiation. Because there are opportunities for activational sex differences to emerge, a comprehensive timeline is needed to capture changes across the rest of postnatal life as well.

Longitudinal *in vivo* Manganese-Enhanced Magnetic Resonance Imaging (MEMRI) in mice addresses many of these shortcomings and is ideally suited for the comprehensive study of neuroanatomical sex differences. This technique builds upon *ex vivo* mouse MRI studies that have been used to capture dimorphisms in the whole brain: from large structures such as the cortex, cerebellum and thalamus; to smaller nuclei in the hypothalamus [18]. In MEMRI, systemic administration of manganese chloride, MnCl_2_, results in visualization of brain architecture, particularly in neonatal neuroanatomy where large-scale changes of cellular composition may pose challenges for optimizing MRI contrast [19]. MEMRI has been used to acquire *in vivo* images of the brain of postnatal rodents as young as 1 day old, and is well tolerated by neonatal rodents through repeated rounds of imaging [20]. Since MEMRI allows for repeated *in vivo* MRI scans of the same animal, neurodevelopmental sexual dimorphisms can be studied beginning in early life and into adulthood.

Here, we used MEMRI to investigate the development of structural sex differences in the mouse brain beginning from early neonatal life. Male and female C57BL/6J mice were scanned longitudinally with MEMRI across 9 postnatal time points, at days 3, 5, 7, 10, 17, 23, 29, 36, and 65. First, known sex differences in the BNST, MeA and MPON were investigated to affirm that MEMRI is a robust technique for detecting sex differences in the brain. Second, linear mixed-effects modelling was used to identify sexually dimorphic areas across the whole brain and characterize their development. Third, *k*-means clustering was used to identify trajectories of coordinated sexually dimorphic development and reveal networks of functionally connected areas.

## 3 Results

### 3.1 MEMRI captures changes in neonatal brain development without adverse effects

The methodology and image analysis techniques employed here are a direct extension of previous work [20], expanding both the throughput and the imaging window to include adult brain development, and applied to both sexes.

To accommodate the small body size of neonatal mice, custom 3D-printed holders were created, which enabled us to routinely position the mouse brain at the centre of the coil for neonatal mice of varying sizes (Figure 1A). Up to 7 mice were scanned in individual saddle coils (Figure 1B).

**Figure 1:**
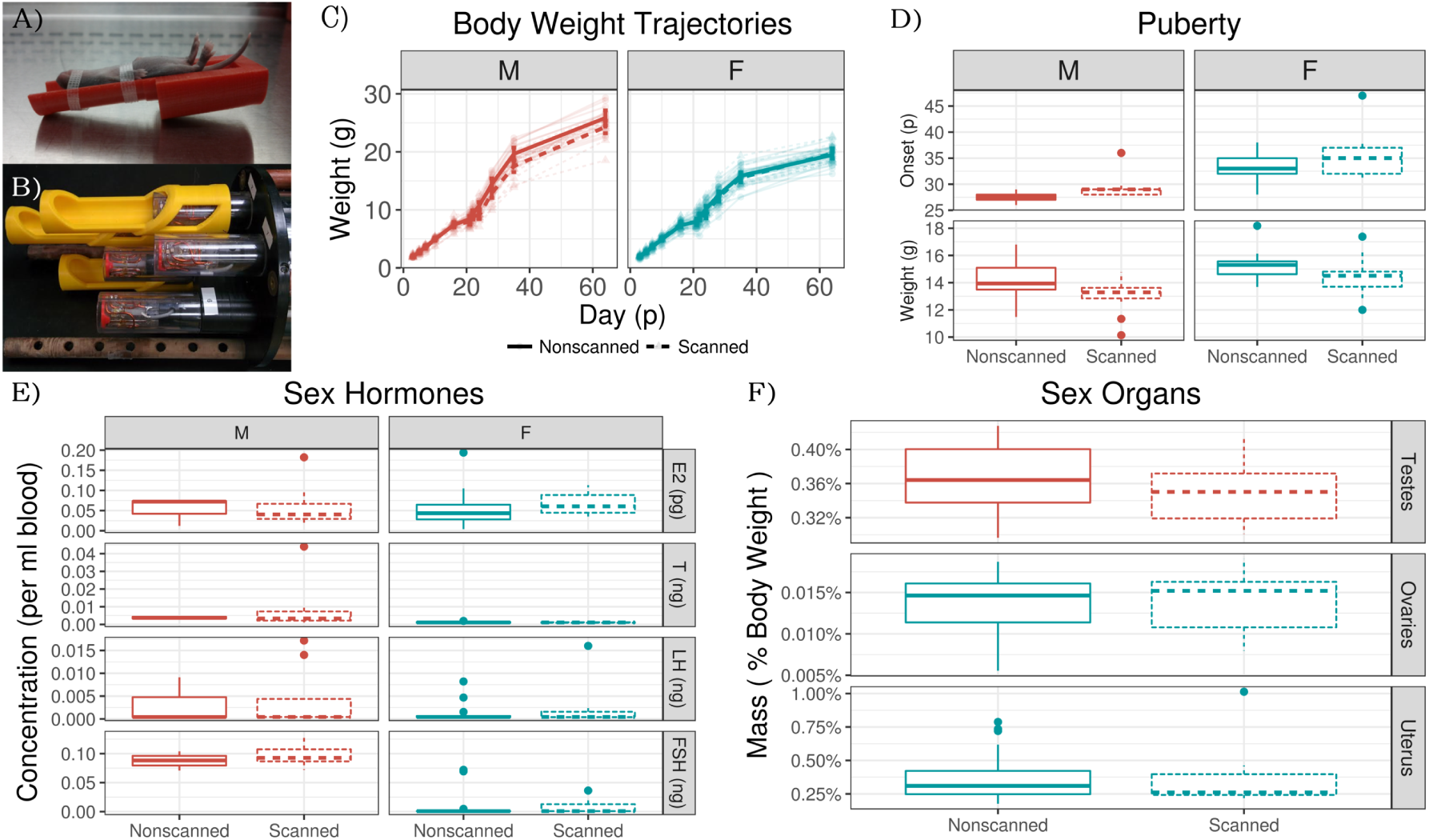
Scanning apparatus, and scanned mice vs. nonscanned mice data. (A) Custom 3D printed holders for neonatal mice. B) Up to seven mice at a time were scanned using a saddle coil array. C) Body weight was not significantly different between scanned and unscanned mice (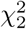 = 0.29,P=0.87). There was also no interaction effect of scanning and sex 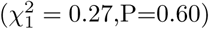. D) Repeated scanning did not have a significant effect on puberty onset (*F*_3,64_ = 24.67,*t*_64_ = 1.56,P=0.12) but did have a significant effect on weight at puberty (*F*_3,64_ = 10.67,*t*_64_ = −2.43,P=0.02). However, neither measure had a sex-scanning interaction (*t*_64_ = −0.045,P=0.96 and *t*_64_ = 0.451,P=0.65). E) Scanning did not have a significant effect on the levels of sex hormones estradiol (*F*_3,41_ = 0.83,*t*_41_ = 1.42,P=0.16), testosterone (*F*_3,41_ = 4.124,*t*_41_ = 0.32,P=0.75), LH (*F*_3,44_ = 0.98,*t*_44_ = 0.56,P=0.58), FSH (*F*_3,32_ = 32.5,*t*_32_ = 0.56,P=0.58). F) Organ weight of testes (*F*_1,18_ = 1.05,*t*_18_ = −1.0,P=0.32), ovaries (*F*_1,28_ = 0.03,*t*_28_ = 0.17,P=0.87), and uteri (*F*_1,28_ = 0.17,*t*_28_ = −0.41,P=0.69) were not affected by repeated scanning

Mice that undergo repeated MEMRI scanning experience increased handling, anaesthesia exposure, and MnCl_2_ injections. To determine if repeated scanning had any effects on development, several measurements were collected from scanned mice and their non-scanned littermates. Weight at puberty (Figure 1D) was significantly affected in scanned mice; however, we did not find a significant effect of scanning on any of the other measurements collected (Figure 1C,E,F), nor did we find evidence of an interaction between scanning and sex. We conclude that the effect of MEMRI scanning on development was small and not sexually biased.

Image registration is necessary for identifying homologous features in MR images, thereby allowing statistical comparison. To accommodate the rapid changes in brain shape during early development (see Supplementary Movie 1), we modified an existing image registration pipeline [21] into a two-level registration pipeline (Figure 2, see Methods 5.4.1 for details). In the first level, images from different mice collected at the same time point were registered together, creating a consensus average brain for each age. In the second level, the age-specific consensus average brains were registered to one another serially to map them all to the p65 consensus average space. By concatenating the transformations from the first and second levels, we achieved point-correspondences between all images and generated a transformation mapping all images — from every mouse and time point — to the p65 consensus average space. Computation of the determinants of these transformations allows measurement of the volumetric differences between all images at individual points in the brain or across defined structures.

**Figure 2:**
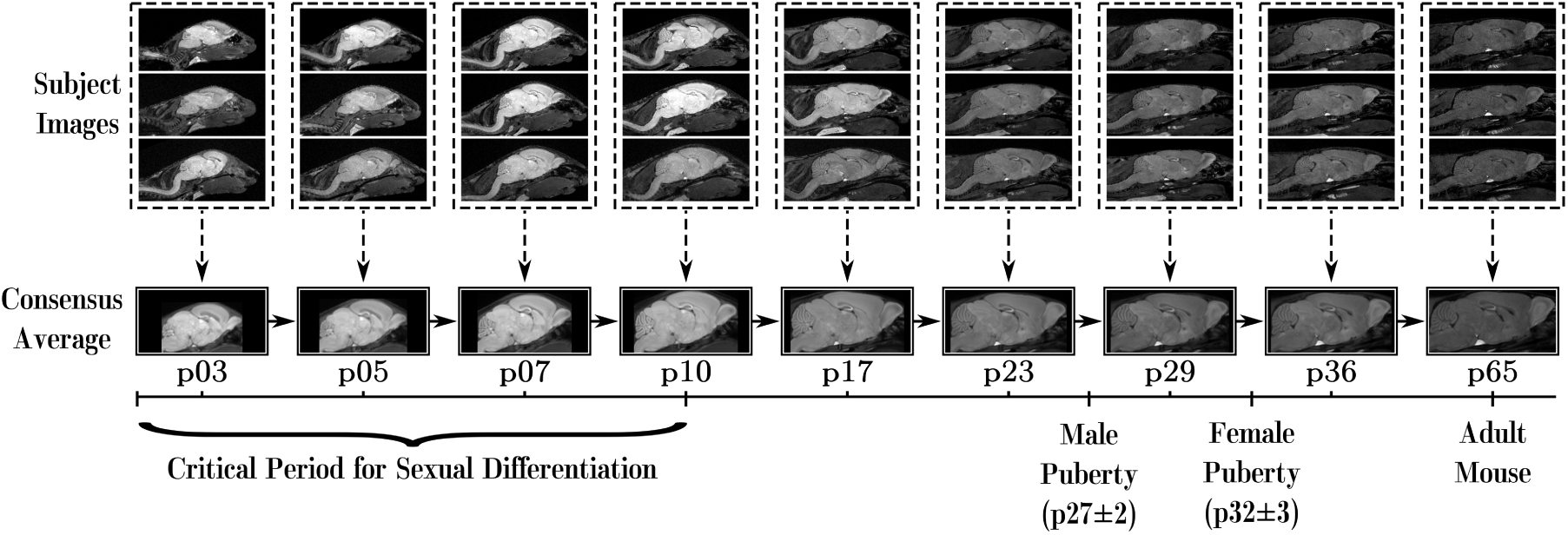
MR images was registered using a two-level approach. In Level 1 (dashed lines), images from each time point were registered together to create a consensus average for each age. In Level 2 (solid lines), the consensus averages for each age were registered to the following adjacent age. The transformations created from concatenating Level 1 (dashed arrows) and Level 2 (solid arrows) allow us to map points in the p65 consensus average to all images.

### 3.2 Canonical and novel sexually dimorphic areas are detectable by MEMRI

To compare structure volumes between sexes, an atlas that segments 161 structures in the adult mouse brain [22] was overlaid onto the p65 consensus average brain. Shown in Figure 3A are segmentations of the MeA, BNST and MPON, which are canonical sexually dimorphic areas. Extensive literature has shown that these sexually dimorphic areas are larger in the male brain. MEMRI captures these sexual dimorphisms (top-panels in Figure 3B-D) and shows that these dimorphisms emerge between p10 and p17. Males tend to have bigger brains than females [11]; as MEMRI allows for whole-brain imaging, we were able to identify dimorphisms in brain structure relative volumes — that is, subjects’ structure volumes divided by their whole-brain volume, in addition to identifying absolute brain volume differences. Like the results based on absolute volumes, males had larger relative volumes of BNST, MeA, and MPON. However, relative volumetric sex differences emerged earlier than the absolute volume differences, around p5. Moreover, several structures, such as the periaqueductal gray (PAG), exhibited no sex-dependent growth differences in absolute volumes, but did exhibit significant differences in relative volumes. Compared to male-enlarged structures, the PAG became relatively larger in females at a later developmental time (around p29). Correction for whole-brain size using relative volume measurements from MEMRI data reveal well-established and novel sexual dimorphisms as well as their time courses, highlighting the strength of whole-brain MRI.

**Figure 3:**
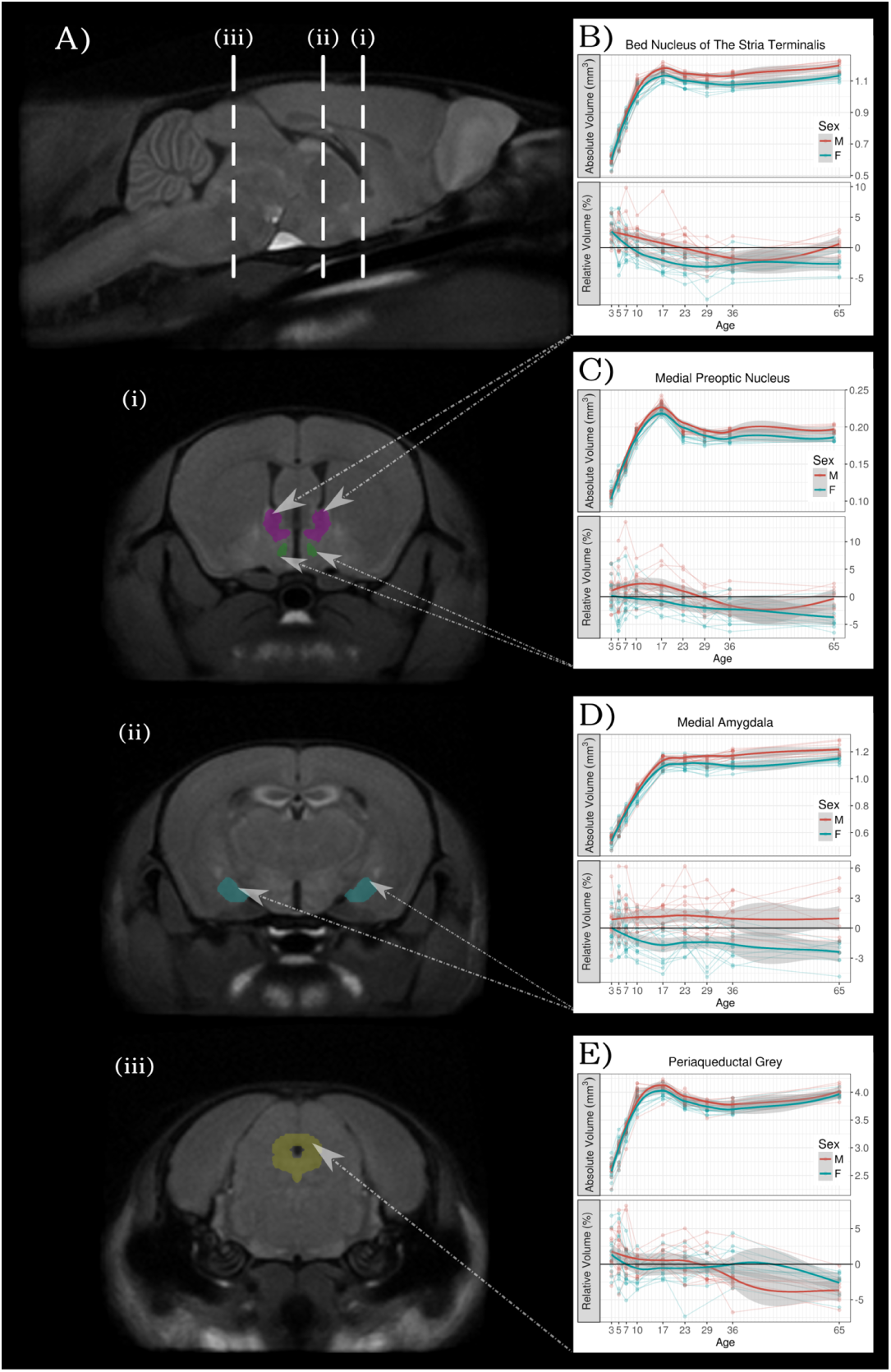
MEMRI captures sex differences in brain structure sizes. A) Sagittal and coronal slices of the average p65 brain showing segmentations of mouse brain structures: Bed Nucleus of the Stria Terminalis (BNST), Medial Preoptic Nucleus (MPON), Medial Amygdala (MeA), and Periaqueductal Grey (PAG). B-E) the absolute and relative volumes of these structures over time. Relative volume corrects for whole-brain size differences between subjects and is expressed as a percent difference from the average volume of the brain structure. Using linear mixed effects models, we recapitulate known canonical sex differences in absolute volumes of the MeA 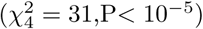, BNST 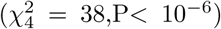, and MPON 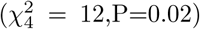. Sex differences in these structures emerge pre-puberty, at around p10. Using relative volumes to correct for whole brain size, we see that sex differences in these structures are preserved but differences emerge very early in development around p5. PAG relative volume also shows a significant effect of interaction between sex and age 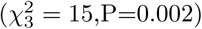, which is not found in the absolute volumes 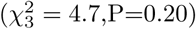. Taken together, these results indicate that relative volumes obtained by MEMRI are a sensitive marker for detecting sexual dimorphism in neuroanatomy.

### 3.3 Relatively larger areas in males emerge in early life, and relatively larger areas in females emerge in post-pubertal life

To investigate the presence of novel sexually dimorphic areas across the whole brain, we examined anatomical differences at the voxel level. The registration pipeline (see Methods 5.4.1) generates a Jacobian determinant field for every image. At every voxel, the value of this field measures the volumetric growth or shrinkage compared to the average brain at p65. Statistics were conducted on the logarithm of the relative Jacobian determinants — that is, the Jacobian determinants after correcting for whole brain size differences.

To test the effect of sex on region volume (see Methods 5.4.4), we ran two linear mixed-effects models on a per-voxel basis: one with fixed effects of sex, age, their interaction and random effect of mouse ID; and a similar one without the effect of sex (and therefore also without sex-age interaction). A log-likelihood test was performed between these two models to assess the significance of sex. Our analysis revealed that sex has widespread influence on neuroanatomical development (Figure 4); including regions such as the cerebellum, midbrain, pons, medulla, PAG, thalamus, hypothalamus, hippocampus, amygdala, caudoputamen, BNST and olfactory bulbs.

**Figure 4:**
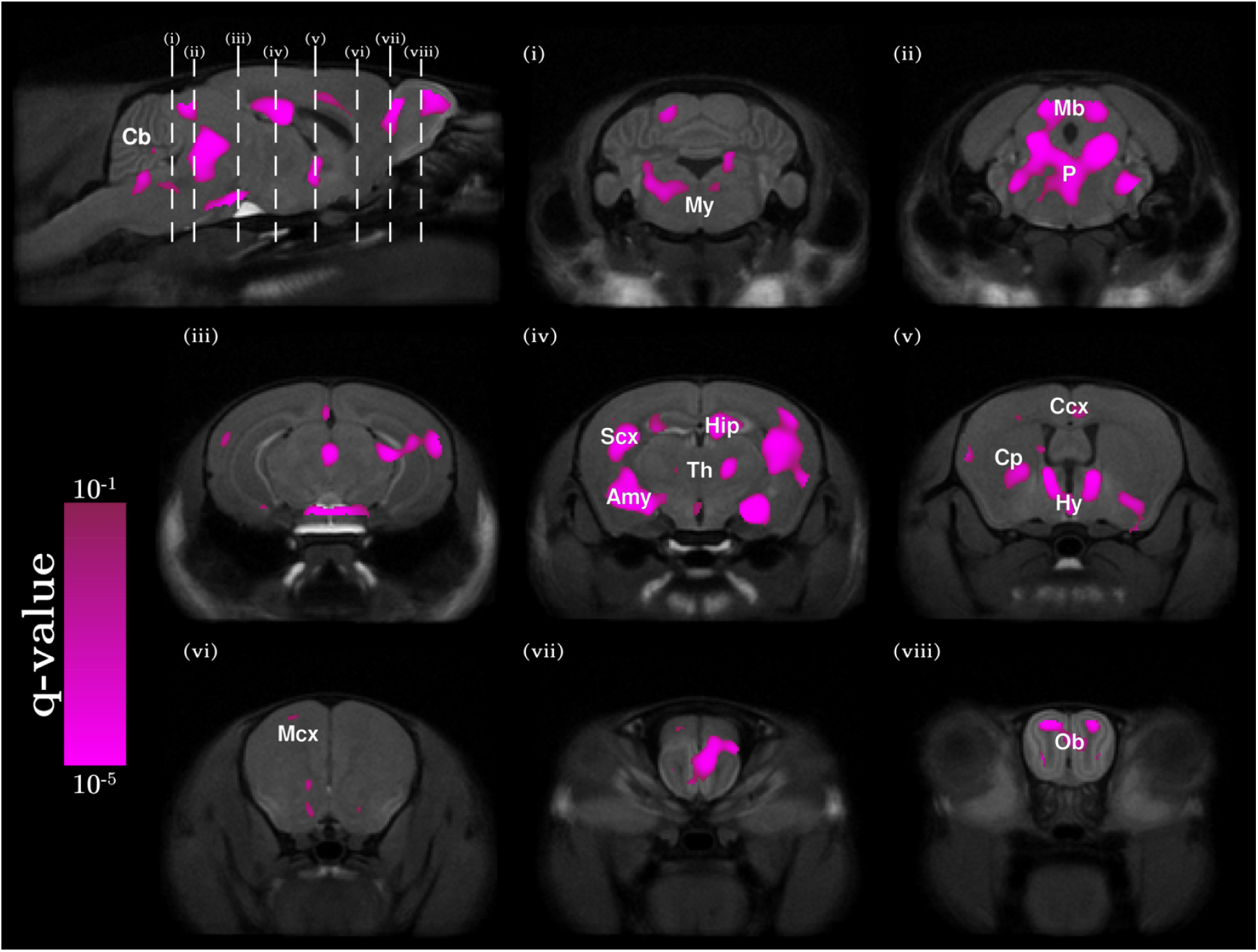
Sexually dimorphic neuroanatomy. Two linear mixed-effects models were fit to the data: Model 1 had sex as a predictor and Model 2 did not. Every voxel was assessed on whether Model 1 had a significantly better fit than Model 2. The resulting q-values (thresholded to q< 0.1) were overlaid on p65 average brain cross-sections, identifying several regions in the brain where volumes are significantly dependent on sex. These sexually dimorphic regions include parts of the cerebellum (Cb), medulla (My), midbrain (Mb), Pons (P), PAG, thalamus (Th), hippocampus (Hip), amygdala (Amy), sensory cortex (SCx), hypothalamus (Hy), cingulate cortex (Ccx), caudoputamen (Cp), BNST, motor cortex (Mcx), and olfactory bulbs (Ob).

To visualize the effect that sex has on brain development at particular time points (see Methods 5.4.5), we fit a model with both age and sex as predictors, but translated the age term such that it was zero at the time point of interest. Nine such models were fit, each centered to one imaging time point. The *t*-statistic field associated with the sex term from each age-centered model was overlaid on the consensus average image from that time point (Figure 5 and Supplementary Movies 2-4). Areas shown in red are relatively larger in males, and areas shown in blue are relatively larger in females. Upon closer analysis of the time course of sexually dimorphic development, it was clear that areas larger in males or larger in females differed in the timing of their emergence. Regions that were relatively larger in males predominated the brain in neonatal, pre-pubertal life and encompassed areas in the MeA, hypothalamus (MPON), BNST, anterior cingulate cortex, hippocampus and olfactory bulbs. In contrast, areas that were relatively larger in females predominated the brain later in life, post-puberty, and encompassed broad, diffuse networks of areas that included parts of the cerebellum, pons, midbrain, PAG, caudoputamen, thalamus and cortex.

**Figure 5:**
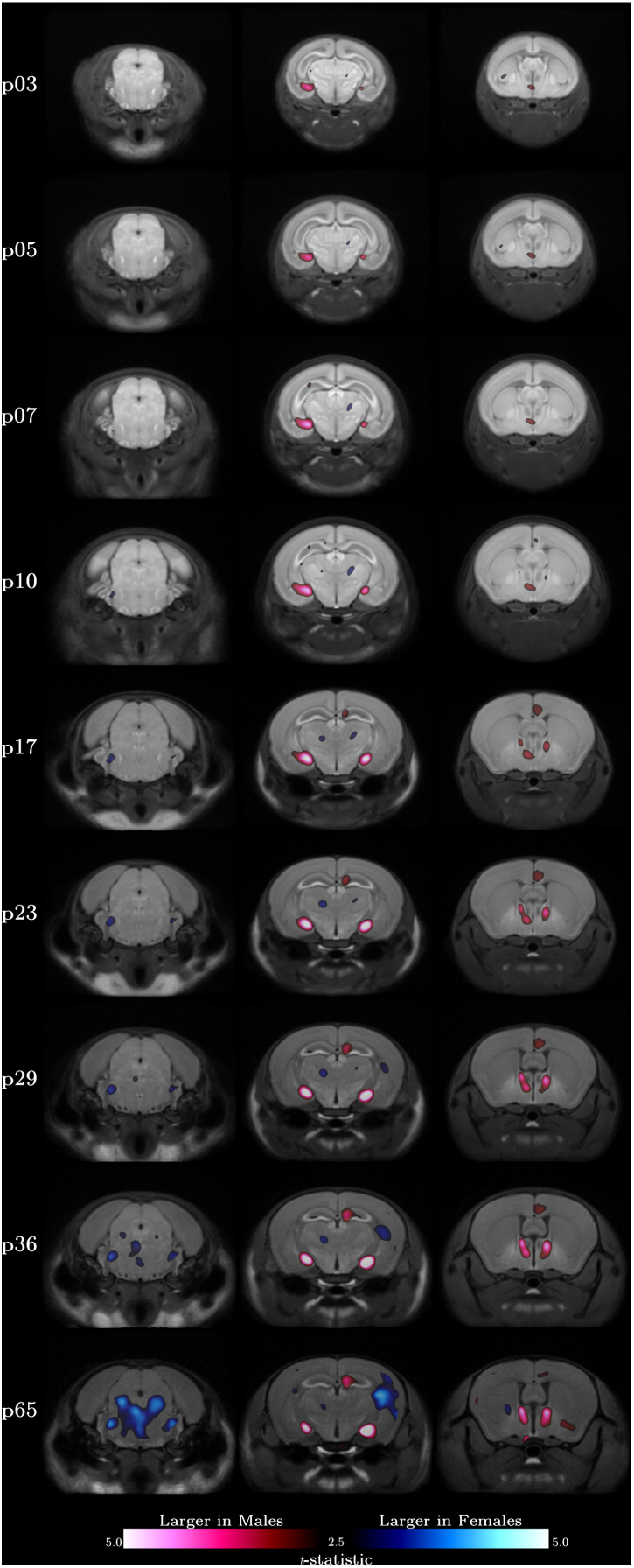
Expansion of neuroanatomical structures in males and females over time. Each column follows a coronal cross-section of the developing brain through the nine experimental time-points with red regions larger in males and blue regions larger in females. Nine age-centered linear-mixed effects models were fit to the data, one for each experimental time point. Each model had identical predictors of sex and age, however the age terms were translated such that they were 0 at the time point of interest. For each time point and corresponding age-centered model, the average brain at the time point was overlaid with the *t*-statistics map associated with the main effect of sex. Statistics are thresholded to 10% FDR in the model centered at p65. Regions relatively larger in males predominate the brain in pre-puberty life (as shown by the second and third image columns). Post-puberty however, regions larger in adult females begin showing dimorphisms.

### 3.4 Clustering sexually dimorphic regions by developmental trajectories reflect functional networks and spatial gene expression patterns

We sought to identify groups of sexually dimorphic regions that had similar developmental trajectories. For all sexually dimorphic voxels identified from our prior analysis (Figure 4), we computed the average male determinant and the average female determinant for each time point. We then took the log ratio of the male and female determinants; this log male-female ratio is positive for regions larger in males and negative for regions larger in females. Using *k*-means, we then clustered the voxels by their log male-female ratio time-series into 4 developmental trajectories (Figure 6). To study differences in growth rate of sexually dimorphic regions, we first fit cubic splines at every voxel for every individual, and differentiated the result to estimate growth rate (see Methods 5.4.6).

**Figure 6:**
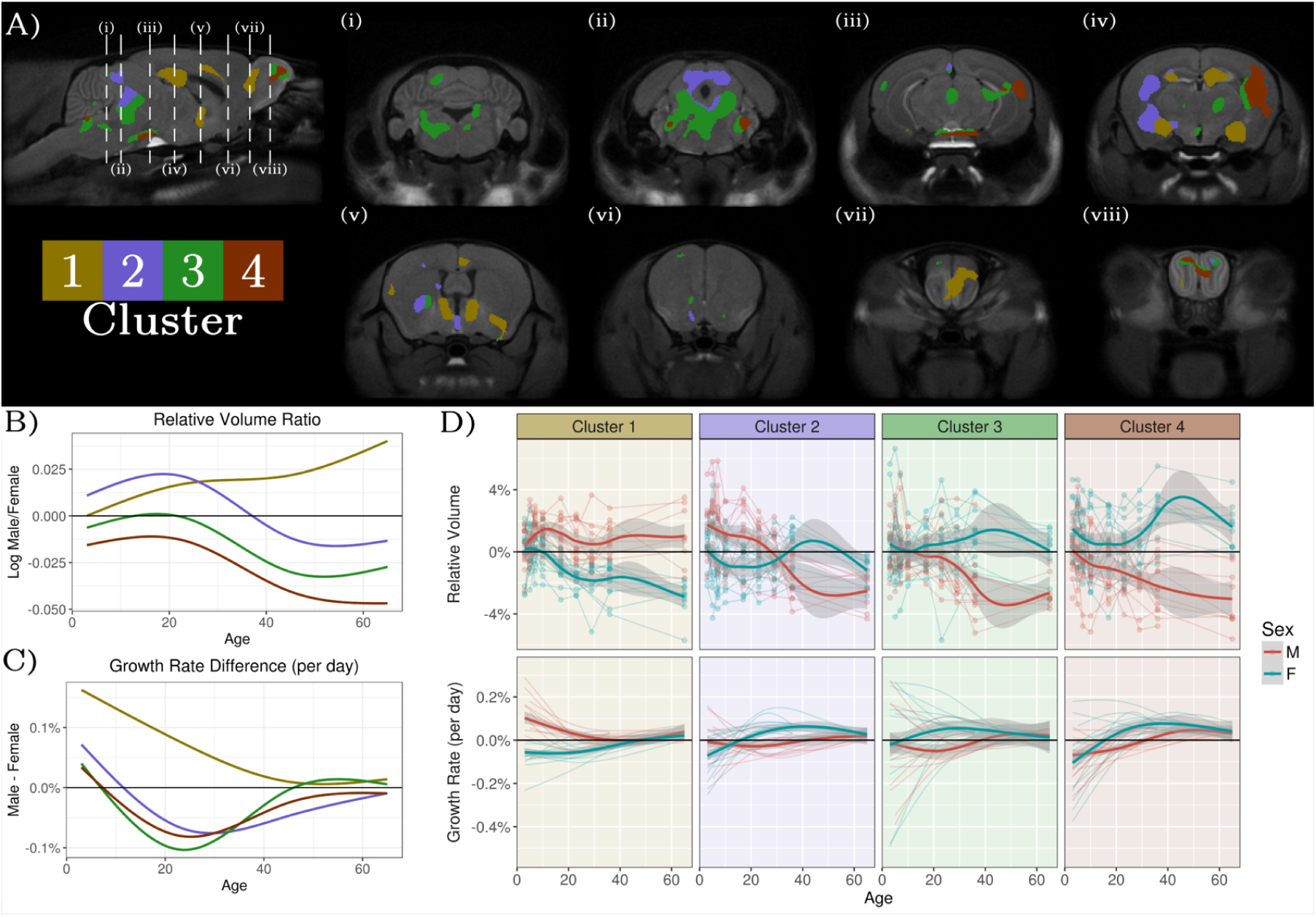
Coordinated growth of sexually dimorphic functional networks. Sexually-dimorphic voxels with similar log ratio of the male determinants to female determinants (log male/female) through time were clustered into 4 groups using *k*-means. A) Results of the clustering analysis on sexually dimorphic voxels. B) Log male/female ratios and C) Male-Female growth rate differences for the different clusters. D) Average volume and growth rate in each cluster for each individual. Cluster 1 corresponds to regions larger in males and this dimorphism emerges early in development. Regions involved in the vomeronasal system, which processes pheromonal information, are found in this cluster. Cluster 2 trajectory switches from being larger in males in early life, to larger in females post-puberty. Parts of the sensory cortex and PAG belong to this cluster. Cluster 3 voxels are not sexually dimorphic in early life and become larger in females over the course of development. This cluster includes association related areas such as parts of the central thalamus and temporal association cortex; and motor related areas such as parts of the hindbrain, cerebellum, caudoputamen, and motor cortex. Finally, Cluster 4 represent voxels that are larger in females throughout development and include parts of the sensory cortex.

Cluster 1 describes areas in the brain that were relatively larger in males throughout development and this dimorphism emerged early in development. Voxels from this cluster reside in the BNST, MeA, MPON, olfactory bulbs, hippocampus, cingulate cortex and pallidum. This cluster had a higher growth rate in males, which stabilized in later life. Clusters 2, 3 and 4 describe areas that became relatively larger in females by adulthood, but were different in early development. Similar to Cluster 1, voxels in Cluster 2 began as relatively larger in males. However, during peripubertal development, they transitioned to becoming relatively larger in females. Voxels in this cluster fall into multiple areas of the brain, including the PAG, superior colliculus, cortex, amygdala and caudoputamen. Cluster 3 showed little sex difference in early life but across puberty, a sex difference emerged as these areas became relatively larger in females. Areas in Cluster 3 reside in the cerebellum, pons, medulla, midbrain, PAG, thalamus, hippocampus, caudoputamen, nucleus acumbens, cortex and parts of the olfactory bulb. Voxels in Cluster 4 were relatively larger in females throughout all of development, and became enhanced following puberty. The majority of voxels in this cluster are located in large parts of the sensory cortices. These clusters had similar patterns of growth rate. Preferential growth in males happened early in life and was followed by a prolonged phase of preferential growth in females around puberty.

Pre-existing studies of mouse neuroanatomy have explored potential underlying biological factors that affect sex differences in the brain. To examine whether our results coincide with any of these factors, we compared our clusters in the developing brain to existing literature regarding spatial patterns of sexual dimorphisms in the adult mouse brain. Sex chromosome aneuploidy studies reveal volumetric decreases in the cerebello-thalamo-cortical system associated with the Y-chromosome [23], which is consistent with parts of Cluster 3. Studies on Four Core Genotype mice revealed that the majority of sexually dimorphic brain regions can be divided into regions associated with sex steroids and regions associated with sex chromosomes [17]. Our developmental clusters did not overlap exclusively with steroid-associated regions or chromosome-associated regions, which suggests that both hormonal and chromosomal effects drive sexually dimorphic development in each of the identified clusters.

We also compared regions of sexually dimorphic development with spatial gene expression data from the Allen Brain Institute [24], which has genome-wide gene expression maps in the adult male mouse brain. We were able to identify genes with preferential spatial expression in sexually dimorphic regions and found that several preferentially expressed genes, such as *Esr2* (Figure 7A) and *Esr1* (Supplementary), are known to be involved with sexualization of the brain (full list of genes in the appendix). Furthermore, the expression patterns of the most preferentially expressed genes we identified, *Slc6a4* (Figure 7B) and *Tph2* (Supplementary), are known to change over the course of development [25]—particularly in the mid- and hind-brain. Compared to a background set of genes from all chromosomes, we also found that sex chromosome genes have a significantly higher likelihood of preferential spatial expression in regions of sexually dimorphic development (Figure 7C).

**Figure 7:**
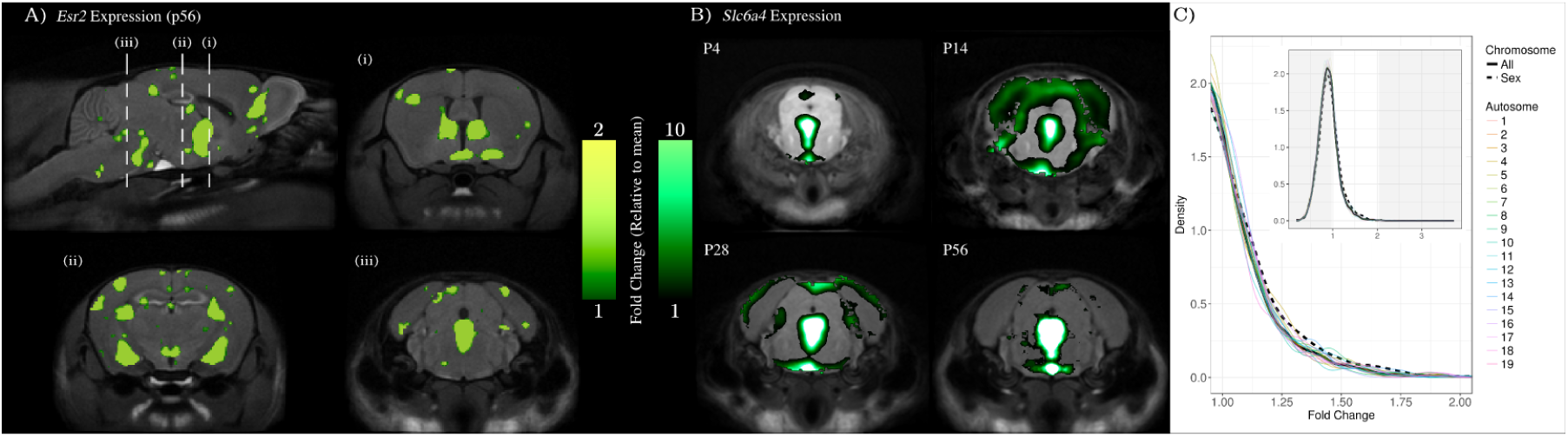
Gene expression patterns overlaid onto MRI results A) Spatial Gene Expression of Estrogen Receptor 2 (*Esr2*) in the adult mouse brain (ABI Dataset ID: 71670737) overlaid onto the MRI average in our study. *Esr2* and *Esr1* genes (expression map in Supplementary) were found to be preferentially expressed in regions with sexually dimorphic development. To measure preferential expression, we used a fold-change measure dividing the mean expression signal in regions with sexually dimorphic development by the mean expression in the brain. *Esr2* had fold-change of 2.9 relative to the whole brain and *Esr1* had a fold-change of 2.1. This was primarily driven by expression in Cluster 1, where the genes had fold-changes of 9.9 and 6.1, respectively. B) *Slc6a4* expression throughout development. *Slc6a4* had the highest preferential expression (fold-change of 3.7) in sexually dimorphic regions primarily driven by expression in Cluster 3 (7.0 fold-change). Timecourse of Slc6a4 gene expression reveals peak expression in mid- and hind-brain between p4 and p28. C) Genes on sex chromosomes have a higher likelihood of preferential spatial expression in regions with sexually dimorphic development. We computed the fold-change for all the genes in the Allen Brain Atlas and created the density plot marked by the solid line. The dashed line indicates the density plot associated with only the genes on the sex chromosomes and the coloured lines represent genes on different chromosomes. In both the overall density plot (inset), and on the zoomed density plot, we see that genes on sex chromosomes have significant preferential expression bias (One-sample Kolmogorov-Smirnov test: *D*^−^ = 0.052, *P* = 0.02, *n* = 730).

## 4 Discussion

Longitudinal MEMRI captures the emergence and development of sex differences in brain anatomy. We recapitulated known sex differences in the brain and identified new regions where development trajectories are influenced by sex. We discovered that differences in neuroanatomical size emerge at different developmental times between males and females: relatively larger areas in the male brain emerge early in development, and relatively larger areas in females emerge peri- and post-puberty. Clustering regions based on shared sexually dimorphic development revealed networks of areas that are functionally connected. Furthermore, by examining spatial gene expression, we found that these brain regions preferentially express genes on sex chromosomes and genes known to be involved with sexual differentiation of the brain.

Examining canonical sexually dimorphic areas with MEMRI largely recapitulates what is known about their development from rodent histology studies. In the BNST, significant differences in cell number and brain volume emerge between postnatal days 9 to 11 [8] following an increase in rates of apoptosis in the female BNST in neonatal life [26]. The MPON sex difference is also due to significantly more cells in the central part of the male MPON, from increased rates of apoptosis in females, emerging by postnatal days 7 to 10 [10] [27]. Part of the MeA sex difference depends on differences in synaptic organization around postnatal day 21 [28], particularly in the middle layer of the medial amygdala [9]. The developmental timing reported for these areas corresponds with our observations that differences emerge in pre-pubertal life. Slight discrepancies in timing of these sexual dimorphisms can likely be attributed to methodological differences, namely the histological study of specific nuclei, layers or areas versus whole structure volume measurements by MRI.

Repeated MEMRI scanning allows for longitudinal observation of the same individual, increasing our sensitivity to detect subtle differences in brain development that emerge across a comprehensive time period. Furthermore, it also allows us to correct for whole-brain size differences to better reveal sexual dimorphisms in neuroanatomy. These aspects of our methodology have enabled us to identify major characteristics about the development of sex differences across the brain. First, developmental periods of relative change in male and female brains are different. Relatively larger areas in males predominate the brain in early, pre-pubertal life and relatively larger areas in females predominate the brain in later post-pubertal life. Similar patterns were also seen without correcting for whole-brain size differences (see supplementary). These pattern mirrors what is known about sex differences in age of onset for many psychiatric disorders: males are more likely to be diagnosed with disorders that are developmental in nature, and have an onset during childhood, while females are disproportionately diagnosed with disorders that have an emotional nature and emerge in adolescence and young adulthood [4]. Periods of relative change in the brain can be considered both as windows of brain development and windows of vulnerability when development goes awry [29]. It is possible, therefore, that periods of relative change in males and females during development are likewise associated with sex-specific periods of vulnerability.

What follows below are examples of developmental periods related to sex that are specific to males and females, and the implications of when processes become aberrant. In neonatal life, males have high levels of circulating testosterone, which becomes aromatized to estradiol; this estradiol plays a crucial role in the sexual differentiation of the brain and modulates many cellular processes [15]. A vulnerability for neurodevelopmental disorders is conferred to males in early life if aberrant estradiol-related action occurs. An example is found in autism: higher rates of autism-like behaviour is linked to fetal testosterone levels [30], and girls with congenital adrenal hyperplasia, who are exposed to elevated gestational testosterone levels, exhibit more autistic traits than unaffected females [31]. Later in life, the presence of hormones affects both the male and female brain [32] [33]; however, in females specifically, this change is linked with estradiol levels [34], since ovarian hormones further feminize the brain [35]. During adolescence, sex differences in many psychiatric disorders emerge [36], with female-biased psychiatric disorders increasing in prevalence. Furthermore, fluctuation of hormone levels during the menstrual cycle, pregnancy and parturition can affect mood and risk of depression [37] [38] [39]. The presence of these hormonal transitional periods in the adult female brain may confer a unique vulnerability for mood and anxiety disorders to women [40]. The ability of MEMRI to detect sex differences in relative neuroanatomical change provides insight on developmental times that may be particularly important for males or females in understanding disorders that show sex bias.

The clustering analysis shows 4 groups of areas, each cluster characterized by a unique trajectory of sexually dimorphic development over time. As log male/female is a size comparison, clustering by this metric allows us to provide insight on areas that cluster together as networks of connected structures that may share function [41]. Most voxels from cluster 1 reside in areas that are well known to be sexually dimorphic and larger in males; they include the BNST, MeA, MPON and parts of the olfactory bulbs [6] [7] [42]. These areas are part of a functionally and structurally connected network related to the vomeronasal system. This system processes pheromonal information to mediate a wide range of social behaviour in both sexes, including fear, reproductive and sexual behaviour, as well as maternal behaviour in females [43]. Each of these structures has been shown to be larger in males than in females, and this sex difference is dependent on neonatal circulating estradiol during the critical period of sexual differentiation [7] [9] [44] [45].

Many structures from the clustering analysis are also structures involved in processes of pain and analgesia. There is robust clinical and laboratory evidence that indicates there are sex differences in pain and analgesia: females have increased pain sensitivity, lower threshold and tolerance for pain, and present more chronic pain problems in the clinic [1]. Structures from the clustering analysis that are involved in pain processes include: the hippocampus, cortex, anterior cingulate cortex, PAG, amygdala, thalamus, caudoputamen, nucleus acumbens, medulla and cerebellum [46] [47] [48]. Interestingly, these structures come from all 4 clusters. Those structures from clusters 1 and 2 (hippocampus, cingulate cortex, PAG, amygdala) are relatively larger in males in early pre-pubertal life, while those structures from clusters 2, 3 and 4 (PAG, amygdala, thalamus, caudoputamen, nucleus accumbens, medulla, cerebellum, cortex) are relatively larger in females in later, post-pubertal life. Indeed, there is evidence suggesting that pain processes are sensitive to, and can be modulated by both neonatal gonadal hormones and activational hormones that are present later in life [49]. In males, certain pain processes can be modulated by manipulating testosterone levels during neonatal development, but not during adulthood, suggesting that areas involved in pain are responsive to hormones during early life [50]. Conversely, in females, both naturally occurring and externally induced changes of hormone levels in adulthood are associated with alterations in pain processes [51] [52], suggesting that the areas involved are influenced during pubertal and post-pubertal development. However, there also exists pain processes which can be modulated by manipulation of sex hormones in both neonatal life and adult life [53]. The pain-related areas from our clustering analyses include regions that are both relatively larger in males in early life, and relatively larger in females in later life, which may be a reflection of their sensitivity to hormones during both neonatal and adult life. This is one example of how understanding the trajectories of male-female differences in brain structure may further our understanding of important differences in function.

Cluster 4 describes areas of the brain that are relatively larger in females in early life, which then become enhanced post-pubertally. Areas in the sensory cortices are featured prominently amongst these areas. In humans, females have greater cortical thickness in many parts of the cortex, particularly in temporal and parietal areas, even without controlling for differences in brain size [54]. This sex difference is present beginning in late childhood, although developmental periods before late childhood have not been examined in humans, and so the precise timing of when this difference emerges is not known. Although we acknowledge differences between mice and humans, this cluster represents areas where new knowledge from longitudinal studies in mice may be able to supplement what is known from human studies.

Overlaying gene expression maps onto our images provides insight into the underlying causes that drive our MRI results. *Esr1* and *Esr2* encode for Estrogen Receptor alpha (ER-*α*) and beta (ER-*β*), respectively. These nuclear estrogen receptors mediate estradiol-induced processes of masculinization and defeminization of the brain and behaviour [15]. Both genes are highly expressed in canonical sexually dimorphic areas, such as the BNST, MeA and MPON; all of which are known to be affected by estradiol.

The *Slc6a4* gene was also highly preferentially expressed in sexually dimorphic areas. This gene encodes the serotonin transporter (SERT), which affects serotonin levels through re-uptake of intrasynaptic serotonin [55]. Both the number of SERT binding sites and cells expressing SERT can be modulated by estradiol [56] [57]. *Slc6a4* gene expression maps onto sexually dimorphic structures, particularly in the hindbrain, which are influenced by estradiol. Expression of *Slc6a4* changes across development, with peak expression occurring between days p4 and p28, and a known peak at p14 (figure 7). This window of postnatal development is too broad to infer the length of peak expression, or whether peak expression is biased towards neonatal or post-pubertal development. However, variants of the *Slc6a4* gene have been implicated in both neurodevelopmental disorders with early onset, and affective disorders with later onset in peri- and post-pubertal development. These include disorders such as autism, attention deficit hyperactivity disorder and Tourette syndrome, as well as depression, anxiety and obsessive compulsive disoder [58] [59].

In summary, we have used longitudinal MEMRI to study anatomical sex differences across the whole mouse brain to characterize both known and novel male-female differences throughout post-natal development. We have shown that MEMRI is a robust method for detecting neuroanatomical sex differences and their timecourses. We found distinct periods of relative developmental change in males and females, where increased male growth predominates in early life and increased female growth predominates in post-pubertal life. By clustering areas in the brain based on shared sexually dimorphic development, we have revealed networks of areas that are functionally connected and mediate sexually dimorphic processes. These findings demonstrate the power of whole-brain *in vivo* MEMRI for examining the development of sex differences, and the importance of studying sex differences across the whole brain across a comprehensive temporal context that begins in neonatal life.

## 5 Methods

### 5.1 Animals and non-imaging procedures

Male and female C57BL/6 mice were scanned longitudinally across 9 postnatal day (p) time points: p3, p5, p7, p10, p17, p23, p29, p36 and p65. Number of mice at each time point are as follows: p3 (n=10 males, n=12 females), p5 (n=14 males, n=14 females), p7 (n=14 males, n=14 females), p10 (n=14 males, n=14 females), p17 (n=14 males, n=14 females), p23 (n=14 males, n=14 females), p29 (n=11 males, n=11 females), p36 (n=14 males, n=14 females), p65 (n=12 males, n=12 females). Each individual mouse was scanned at all time points; discrepancies in mouse number at some time points was due to occasional scanner issues that caused certain scans at time points to be excluded. Number of mouse pups in each litter was reduced to 6 to ensure equal manganese intake by pups through maternal milk. Because manganese is administered to neonatal mice through maternal milk, non-scanned littermates were also exposed to manganese for the first 10 days of life. The ratio of male to female mice in each litter was kept equal. To differentiate neonatal pups, mice received black ink tattoos on their paws at p2 (AIMS Lab Animal Tattoo Kit, AT-3 General Rodent Tattoo System).

Two pups from each cage were chosen for longitudinal scanning. Both scanned and nonscanned littermates were weighed either on the day of scanning, or the day prior to scanning as a measure of overall growth throughout the experiment. Mice were weaned at p21 and separated into cages by sex. Post-weaning, mice were assessed for puberty daily. First occurrence of preputial separation (PS) after weaning was used as an indicator of puberty for male mice [60], and first occurrence of vaginal opening (VO) after weaning was used as an indicator of puberty for females [61]. Weight at puberty was also recorded. Mice were housed in cages with up to 4 mice, and maintained on a 12-hour light/dark cycle, with ad libitum access to food and water.

At postnatal day 66, blood was collected for hormone level measurements and organs (gonads and uteri) were dissected out of scanned and nonscanned mice, to be weighed. Mice were anaesthetized with 1-4% isoflurane in air. While under anaesthesia, blood for plasma was collected via cardiac perfusion by opening the thoracic cavity and drawing blood from the left ventricle. Mice then underwent cervical dislocation, and ovaries and uteri were dissected from female mice, while testes were dissected from male mice. Dissected tissues were placed in a dish with phosphate-bu ered saline and excess fat was removed from the tissues under a light microscope. Before weighing, tissues were blotted on a Kimwipe to remove excess liquid. Tissues were weighed on an analytical scale accurate to 0.1 mg. Blood samples were sent to The Endocrine Technologies Support Core at the Oregon National Primate Research Center (Beaverton, OR). Barring samples of insu cient size, all samples were analysed for estradiol, testosterone, follicle-stimulating hormone (FSH) and luteinizing hormone (LH). All experiments were approved by The Centre for Phenogenomics Animal Care Committee.

### 5.2 *In vivo* imaging

Up to 7 mice of the same age were scanned simultaneously *in vivo*. 24 hours prior to the scan, mice received a 0.4 mmol/kg dose of 30 mM manganese chloride (MnCl_2_) solution. For mice 10 days and younger (neonates), MnCl_2_ was provided through maternal milk by injecting mothers 24 hours prior to the scan. Mice 17 days and older received intraperitoneal injections directly 24 hours prior to the scan. Throughout the scan, bore temperature was maintained at 29 °C, and a steady stream of 1-2% isoflurane was used to keep the mice anaesthetized. Respiration was monitored throughout the scan. Respiratory pillows were used for mice 17 days and older; self-gated signals from a modified 3D gradient-echo sequence [62] provided respiratory motion information for neonatal mice which were too small for respiratory pillow use.

A multi-channel, 7.0 Tesla, 40 cm diameter bore magnet MRI scanner (Varian Inc. Palo Alto, CA) was used to acquire images of mouse brains. Parameters of the scan are as follows: T1-weighted, 3-D gradient echo sequence, TR = 26 ms, TE = 5.37 ms, flip angle = 37 °, field-of-view = 77 × 20 × 20 mm, matrix size = 854 × 224 × 224, number of averages = 5, total acquisition time = 1 hour and 40 minutes, isotropic resolution = 90 *μ*m. Post-scanning, mice were transferred to a heated cage for 5-10 minutes in order to recover from anaesthesia, and then returned to their home cages.

### 5.3 Analysis of scanned and non-scanned data

Growth was compared between scanned and nonscanned animals by running two linear mixed effects models: both models had fixed effects of sex, age (approximated as a cubic spline), and their interaction, with a random effect of growth for each mouse. One model had an additional fixed effect of type (scanned or nonscanned) and type-sex interaction. The two models were then compared with a likelihood ratio test to assess whether scanning affected growth of mice or if the effect had significant sex-bias. Differences between scanned and nonscanned mice in weight and puberty onset, hormone levels, and organ weights were analysed using linear models.

### 5.4 Image analysis

#### 5.4.1 Longitudinal registration

Image registration allows quantification of anatomical differences between images. For a group of images, this procedure results in a transformation that maps every point in one image to corresponding points in the other images. Thus, the differences between the images are captured by this transformation. Our procedure for image registration is composed of a linear registration, followed by a series of non-linear registrations. The linear registration applies global translation, rotation, scaling, and shearing to align images. Information regarding global deformations (i.e. the overall brain sizes) are stored in these transformation models. Linear registration was performed using the mni autoreg tools [63]. The non-linear registration was performed using the ANTs toolkit [64] and creates a vector field that maps every point in one image to another. This transformation model provides information about localized deformation. The Pydpiper toolkit extends the processes described above to group-wise registrations (described in detail by [21]). Pydpiper takes multiple images as inputs, and outputs a consensus average; as well as linear and non-linear transforms that map the consensus average to all input images.

We modified the registration process to accommodate longitudinal data using a two-level approach. In Level 1, group-wise registration was performed on each age. For example; all the p3 brain images were registered together to create a p3 brain average, all the p5 brain images registered together to create a p5 average, etc. The results of this level are consensus averages of each age and their appropriate transforms to the input images, however the results do not capture deformations across time. Time-dependent deformations are captured by Level 2 of the registration, where the consensus average from each time point is registered to the average from the following time point (p3 average registered to p5 average, p5 average to p7 average, etc). The final step in the registration is to concatenate the transforms from both levels so all images can be mapped to the p65 consensus average brain. For example, to align the image of a p29 subject brain to the p65 average brain, the following transformations are applied: p29 subject to p29 average, p29 average to p36 average, p36 average to P65 average, where the first transformation is obtained in Level 1 and the remaining from Level 2.

The two-level registration procedure creates transforms that map the p65 consensus average to every image. As described earlier, each transform contains a global transformation (derived from the linear registration) and local transformations (derived from the non-linear registration). We used deformation-based morphometry to analyze these transformations. First, the transformation vector field is converted into a Jacobian determinant scalar field. Each point in consensus average has a scalar value associated with it characterizing the degree to which volume elements (voxels) had to grow or shrink to map to the individual images. Thus, the volumetric differences between images are captured by the Jacobian determinants. Determinants of the total transformations (global+local) are called absolute Jacobian determinants as they characterize the true volumetric differences between the images and the p65 consensus average. Relative Jacobian determinants are the determinants of only the local transformations and characterize volumetric differences with the overall effect of brain size removed. The advantage of relative Jacobians is that they can eliminate variability due to overall size, and can reveal relative neuroanatomical differences otherwise difficult to detect. We took the logarithm of absolute and relative Jacobians determinants prior to statistical analysis. Regions with negative log determinants suggest that the region is smaller than the consensus average, while regions with positive log determinants suggests that the region is larger.

To perform volume analysis on structures, we registered an MRI-atlas [22] onto the p65 average. Since subject images from all ages were registered to the p65 average, structures outlined by the p65-registered MRI-atlas could be overlaid onto any image, enabling automated quantification of structure volumes over time. We obtained PAG and BNST segmentations from the MRI-atlas [22]; MPON and MeA segmentations were obtained from a modified atlas in which these two structures were manually segmented.

#### 5.4.2 Statistical Analysis

Statistical analysis was performed using linear mixed-effects models using the 1me4 package [65]. By incorporating fixed and random effects, these models are appropriate for data from the same subject over time and enable more powerful analysis of longitudinal studies. The model formula is given below [66]:

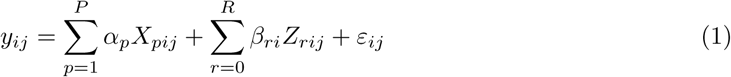

In (1), for a particular mouse *i* measured at a particular time point *j, y_ij_* is the response variable we want to model, *P* is the total number of fixed effects, the matrix *X* represents our fixed effects with *X_pij_* and *α_p_* being the value of the *p*th fixed effect and its coefficient, *R* is the total number of random effects, the matrix *Z* represents our random effects with *Z_rij_* and *β_ri_* representing the value of the *r*th random effect for the *i*th mouse and its coefficient, and *ε_ij_* represent the residuals assumed to be independent and normally distributed.

The response variable *y_ij_* can represent any volumetric measurement, and the predictors are flexible enough to handle the various analysis we performed. When analyzing structures, *y_ij_* represents the structure volume; and when analyzing voxels, *y_ij_* represents the relative log determinant at that voxel.

To perform significance testing for a set of *q* effects (i.e. *q* = {*p*_1_, *p*_2_, …, *p_Q_*}), we fit the data with both the full model (1) and a similar model without the particular effects:

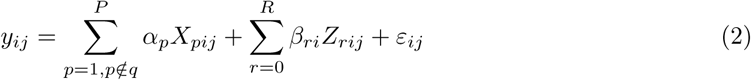

We used the standard likelihood-ratio test to assess whether the full model (1) fits the data significantly better than the partial model (2). Given data, the test statistic *D* can be computed from the maximum likelihood of the full model *L_f_* and the partial model *L_p_*:

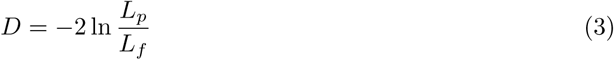

Equation (3) follows the *χ*^2^ distributions with degrees-of-freedom being equal *Q*—the difference between the number of parameters in the full model and the partial model. We can thus compute p-values to measure the significance of the *q* effects. Finally, we used false discovery rate [67] to correct for multiple comparisons.

#### 5.4.3 Sexual Dimorphisms in Canonical Structures

To perform volume analysis on structures, we registered an MRI-atlas onto the p65 average. Since subject images from all ages were registered to the p65 average, structures outlined by the p65-registered MRI-atlas can be overlaid onto any image, enabling automated quantification of structure volumes over time. To test the significance of sex on the structure volumes, we fit two models. Model 1 contains fixed effects sex *s* and age *t*, as well as interaction terms. Growth was modeled as a function of age using natural cubic splines. These splines have three basis functions and the *k*th basis functions is represented by *f_k_*(*t*). Model 1 also had a random intercept for each individual mouse *β_i_*. Based on the general equation (1), the formula for Model 1 is given below:

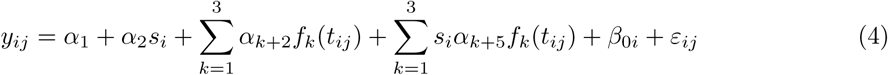

Model 2 was identical to Model 1 but with no effect of sex and no interaction terms. The formula is:

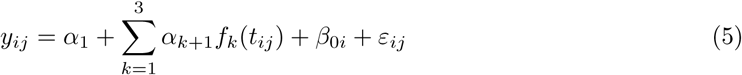

The significance of sex can be computed from the likelihood ratio of the two models. To ensure results were not due to the growth models chosen for fixed and random effects, we experimented with different fixed and random effects. We changed the growth models in fixed effects to natural quadratic and quartic splines. We also imparted more complex random effects structure by providing each individual mouse with random growth trajectories of order 2 natural splines (models with orders beyond 2 failed convergence). These alternative models yielded results that were not substantively different from the ones presented.

#### 5.4.4 Sexual Dimorphisms in Voxels

The effect of sex on voxel determinants was analysed in a similar way by first fitting two models for every voxel in the brain. Model 1 predicted the relative log determinants at that voxel (determinants after correcting for different whole-brain sizes) using fixed effects of sex, age, and their interaction and random intercept for each mouse. Growth was modelled as a linear function of age.

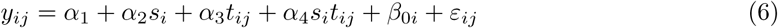

Model 2 was identical to Model 1 but with no effect of sex and no interaction between sex and age.

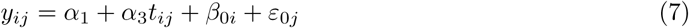

Likelihood-ratio statistic was computed for every voxel comparing the fit between Model 1 and Model 2 and this statistic was used to compute the significance of sex. We experimented with different growth models—growth modelled as a linear and quadratic function of age and random effect of growth for each individual mouse *β*_1*i*_*t*_*ij*_—and observed similar regions of the brain exhibiting sexual dimorphism.

#### 5.4.5 Age-Centered models

To visualize sexual dimorphisms at a particular age *t′*, we used an age-centered model. Similar to the model in (6), the age-centered model references all ages to *t′*, so that the time-independent fixed sex effect (*α*_2_) represents the age-specific difference.

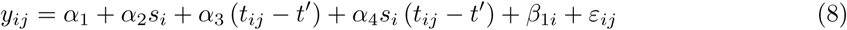

For each voxel, we then extracted the coefficients associated with sex at this age of interest and assigned significance values to the sex effect using the Satterthwaite approximation [68].

#### 5.4.6 Sexual Dimorphism Clusters and Gene Expression

To examine developmental patterns amongst sexually dimorphic areas, *k*-means clustering [69] was used to find groups of voxels that show the same pattern of sexual dimorphism across time. Sexual dimorphism was defined by log male/female of the relative Jacobian determinant. Voxels that showed significance of sex at a false discovery rate of 10%, from the linear mixed effects modelling analysis were included. Using the Elbow method [70], it was determined that 4 clusters was an appropriate number of clusters for this data.

Growth rate was estimated by fitting the relative determinant of each individual and at each voxel with natural cubic spline functions of age, then differentiating the fitted function with respect to age. We found similar results after fitting data with natural quartic splines instead of cubic. However, linear and quadratic splines did not maintain enough complexity after differentiation to fit growth rate patterns.

We used the Allen Brain Institute’s gene expression dataset to identify genes spatially enriched in our clusters. Although some genes had multiple expression datasets, we only chose the datasets with the least number of missing data voxels for analysis. We also excluded experiments where expression data spanned less than 20% of the brain. Preferential expression of a gene in a cluster was measured using a fold-change measure: mean expression signal in the cluster divided by mean expression signal in the brain.

## 6 Acknowledgements

This work was supported by funding from the Canadian Institute of Health Research and the Ontario Brain Institute. LRQ was supported by a Restracomp Fellowship funded by the Hospital for Sick Children. DJF was supported by an Ontario Graduate Scholarship and a Doctoral Postgraduate Scholarship from the Natural Sciences and Engineering Research Council of Canada. KUS and DHT were supported by NIH grant R01NS038461. The authors wish to thank Sharon Portnoy for MRI sequence development assistance; Christina Corre and Ariane Metcalfe for colony management and technical assistance; Matthijs van Eede, Benjamin Darwin and Chris Hammil for help with computation pipelines.

## 7 Author Contributions

LRQ designed and performed experiments, analysed data, wrote manuscript. DJF analysed data, created figures, wrote manuscript. KUS and DHT provided instruction on neonatal imaging. JD designed and built holders and scanning array. BJN provided guidance with MRI pulse sequence development and image reconstruction. MRP and JPL designed experiments and supervised project. All authors edited manuscript.

